# Statistics of correlations in nonlinear recurrent neural networks

**DOI:** 10.1101/2025.10.27.684819

**Authors:** Germán Mato, Facundo Rigatuso, Gonzalo Torroba

**Affiliations:** Instituto Balseiro, UNCuyo and CNEA, S.C. de Bariloche, Río Negro, R8402AGP, Argentina; Centro Atómico Bariloche and CONICET, S.C. de Bariloche, Río Negro, R8402AGP, Argentina

**Keywords:** neural population dynamics, recurrent neural network, nonlinear dynamics

## Abstract

The statistics of correlations are central quantities characterizing the collective dynamics of recurrent neural networks. We derive exact expressions for the statistics of correlations of nonlinear recurrent networks in the limit of a large number *N* of neurons, including systematic 1*/N* corrections. Our approach uses a path-integral representation of the network’s stochastic dynamics, which reduces the description to a few collective variables and enables efficient computation. This generalizes previous results on linear networks to include a wide family of nonlinear activation functions, which enter as interaction terms in the path integral. These interactions can resolve the instability of the linear theory and yield a strictly positive participation dimension. We present explicit results for power-law activations, revealing scaling behavior controlled by the network coupling. In addition, we introduce a class of activation functions based on Padé approximants and provide analytic predictions for their correlation statistics. Numerical simulations confirm our theoretical results with excellent agreement.

## 1 Introduction

Correlations of neural activity are one of the main tools we have to study the structure and function of the nervous system. They provide crucial information by measuring the statistical interdependencies between the activity of different neurons or neural regions over time [1]. The interpretation of the correlations is not easy because they are not only controlled by direct interactions between the neurons but also by the global dynamical state of the system. For instance, theoretical analysis reveal that if the network is in an asynchronous state the cross-correlations are smaller than the autocorrelations by a factor 1*/N, N* being the network size [2]. This regime has been found to be present in a wide variety of systems [3, 4, 5, 6] under quite general conditions.

Small cross-correlations are indeed found in cortical recordings both in spontaneous and active states [7, 8]. Even if the cross-correlations are small it has been recognized that they can be important for the coding and information transmission properties of networks [6]. For instance, in a study where correlated firing in MT (Middle Temporal area) was measured, it was found that spike counts from adjacent neurons were noisy and only weakly correlated but that even this small amount of correlated noise placed substantial limits on the benefits of signal averaging across a pool [9]. It has been also proposed that weak correlations can have a crucial effect in controlling neural variability. In [10] it is shown that physiological levels of synchrony are sufficient to generate irregular responses found in vivo.

Another important effect of small cross-correlations can be found in the control of the effective dimension of the neural dynamics. Dimensionality is a crucial aspect of the network functionality. As has been observed in [11] high-dimensional representations allow a simple linear readout to generate a large number of different potential responses. In contrast, neural representations based on highly specialized neurons are low dimensional and they preclude a linear readout but display better generalization capabilities. One of the most common tools for quantifying the dimension is the *participation dimension* [12]. This quantity, which is defined in terms of the eigenvalues of the covariance matrix, is a natural continuous measure of dimensionality. It can also be expressed in terms of the first and second moments of the diagonal and non diagonal terms of the covariance matrix. It can be easily seen that even if the cross-correlations are order 1*/N* they have a crucial effect on the participation dimension in the large *N* limit because there are order *N* inputs for each neuron. In fact the leading contribution comes from the relative dispersions of cross-covariances across neurons [13].

This implies that a theoretical understanding of the cross-correlations is necessary even if they scale like 1*/N* in the large *N* limit. The calculation of the relevant moments of the covariance matrix was performed in [14] for random connectivity matrices and linear dynamics. In this work the moments were evaluated using a path-integral representation up to order 1*/N*, allowing to evaluate the participation dimension. The same final result was also derived in [15] using techniques of random matrices. One limitation of these results is that in the linear regime the dynamics becomes unstable when the variance of the weights is too strong. This instability can be suppressed in more realistic systems if sub-linear input – output transfer functions are introduced.

In this paper we present a theoretical analysis of a recurrent neural networks with non-linear transfer functions using path-integral methods. This will lead to a new and conceptually simple representation of the neural correlations in terms of a few collective variables. It will allow us to compute the moments of the covariance matrix including the corrections of order 1*/N* necessary to evaluate the participation dimension. This approach can also be extended to compute all higher order neural correlators, as well as to systematically include 1 subleading 1*/N* corrections. Nonlinear recurrent networks have also been studied with other methods, such as dynamical mean field theory (DMFT) (see for instance [16] and references therein), and random matrix theory [17]. In fact, as we were finishing our work, the preprint [18] appeared. It uses a DMFT ansatz with random matrix techniques to study correlations of the covariance matrix of nonlinear networks. We will compare our results to theirs, finding agreement when they overlap.

The work is organized as follows. In Sec. 2 we introduce the model, the path integral description and the collective fields. In Sec. 3 we evaluate the large *N* limit via a saddle point approximation. This section contains our main result: the generating function for the connected correlations of the network. This can be used to derive all correlation functions; we give explicitly the correlators for the covariance matrix. Using this, we derive an explicit and general expression for the dimension of participation. In this section we also comment on the relation with other approaches.

In Sec. 4 we present various applications of our approach. First we consider power-law activation functions, and derive analytic results for the covariance matrix statistics. Then we introduce a class of “Padé activation” functions, which have various useful theoretical properties. We compute the analytic predictions and perform detailed comparisons with numerical simulations, finding excellent agreement. We discuss 1*/N* corrections and non-equilibrium effects. Finally, Sec. 5 is devoted to the conclusions and future directions. While the main text focuses on the correlations of neural outputs, for completeness in Appendix A we show how to derive the correlations for the inputs as well.

## 2 Model and path integral representation

A Recurrent Neural Network (RNN) for *N* neurons with signal variable *ϕ*_*i*_, *i* = 1, …, *N*, and nonlinear activation function *f* (*ϕ*), is described by the equation

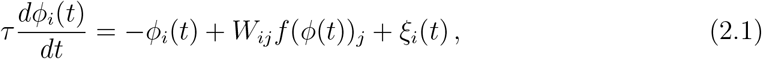

where *ϕ* describes the input neural current, and *f* (*ϕ*), the activation function, gives the neural outputs. We take the convention that all repeated indices are summed up. The number of neurons *i* = 1, …, *N* is taken to be large. We assume that the nonlinear function *f* is regular, odd under *ϕ* → −*ϕ* and vanishes at the origin,

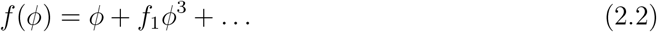

This allows to expand the dynamics of *ϕ* around the fixed point *ϕ* = 0. The coefficient of the linear term can be set to one by a rescaling of *W*. The following results can be generalized to activation functions without parity and cases where *ϕ* acquires an expectation value. But we will restrict our analysis to this simpler case.

The internal noise is a random Gaussian variable with vanishing one-point function and

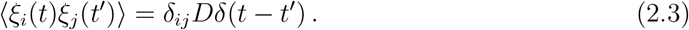

The random connectivity matrix usually fluctuates on time-scales which are much larger than typical single-neuron time variations. So we will take it as a quenched disorder variable described by a Gaussian noise with vanishing expectation value and

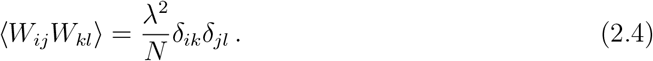

The factor of *N* in this variance is chosen to have a well-defined large *N* limit.

Our goal is to compute the correlation functions of the network outputs,

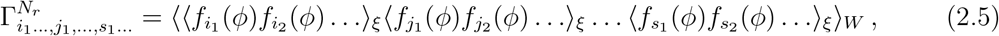

where we kept the insertion times implicit. Here ⟨ … ⟩_*ξ*_ denotes the average over *ξ*, ⟨ … ⟩_*W*_ is the average over *ξ*, and *N*_*r*_ is the number of *ξ*-averages inside the ⟨ … ⟩_*W*_. The calculation of the correlations of the network inputs *ϕ*_*i*_ can be done along similar lines; this is presented in the Appendix A.

It is important to notice that experimental evaluations of the correlations do not involve an average over the coupling variables but empirical averages over some sets of neurons. Here we are assuming that in the limit of large *N* the macroscopic state does not depend on the precise value of each of their random parameters but rather only on their statistics [19]. In other words we assume the system is *self-averaging*. This assumption allows us to use the average over *W* to predict the empirical averages.

Consider first a single *ξ*-average. By parity, correlation functions with an odd number of insertions vanish. Correlation functions with an even number of *f*-insertions can be obtained from the partition function with a source *J*_*ij*_(*t*_1_, *t*_2_) for *f*_*i*_(*ϕ*(*t*_1_))*f*_*j*_(*ϕ*(*t*_2_)). The time dependence of the source allows to compute correlations at different times; however, while for now we will write a partition function for the general case, the main focus of this work will be on time independent (zero frequency) correlations. Introducing a Lagrange multiplier field 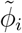 that imposes the stochastic equation (2.1), the path-integral representation for the partition function including the *ξ*-noise average is

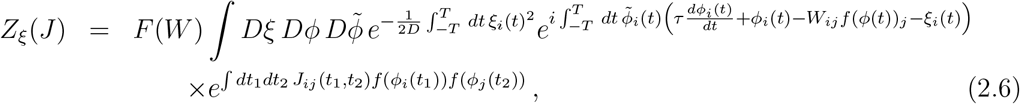

where *F* (*W*) is a normalization factor such that *Z*_*ξ*_(*J* = 0) = 1. In the linear model, a short calculation gives *F* (*W*) as a determinant of the *ϕ* and 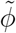 Green’s function, but more generally it is a complicated function that depends on the activation function and cannot be computed in closed form. We will see shortly that its effects are subleading for a large network.

Now let’s turn to the generating function of *W*-averages of the form (2.5). Each *ξ*-average is represented by a generating function (2.6), so we need to introduce *N*_*r*_ replicas; the replica index will be denoted by *a, b*, … = 1, …, *N*_*r*_, while letters *i, j* … are the physical neuron indices. Then the fields in (2.6) acquire two indices: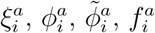, while *W*_*ij*_ has no replica indices because we are considering a single *W*-average. Furthermore, note that by parity only correlation functions of 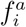 with an even number of *i*-indices are nonvanishing. In order to generate these, it is sufficient to use a bi-source 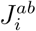 that is diagonal in physical space and nondiagonal in replica space. For instance,

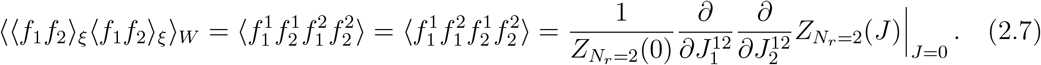

Therefore, the generating function for the statistics of *N*_*r*_ output correlations is

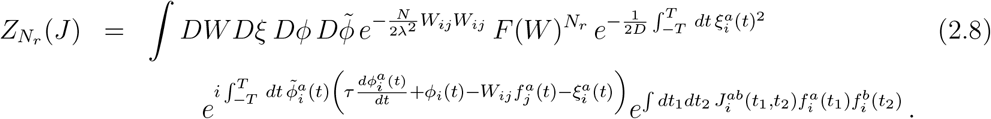

The external source 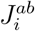 is chosen to be an upper-triangular matrix in replica space.^1^ As long as *N*_*r*_ ≪ *N*, the factor of 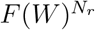 gives a small correction to the Gaussian dynamics of *W*, and so we will neglect it in what follows.

The partition function can be written as a path integral of *e*^−*S*^, with action

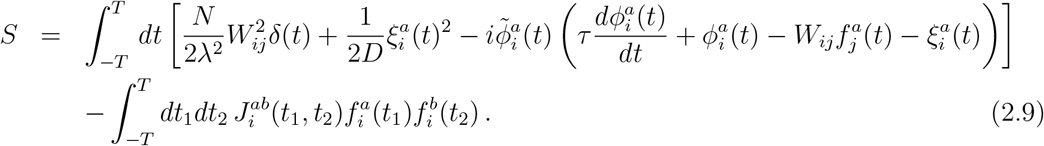

In this paper we will focus for simplicity on equilibrium or zero-frequency correlation functions. It would be very interesting to extend the analysis below to include the time-dependent dynamics of the network, something that we plan to address in future work. In this case we can drop all the time dependence and the time integrals, the path integrals become ordinary integrals, and the action is

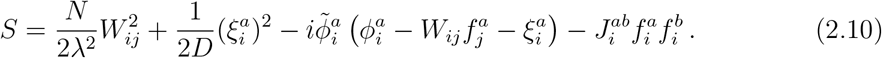

Performing the Gaussian *ξ* and *W*-integrals gives

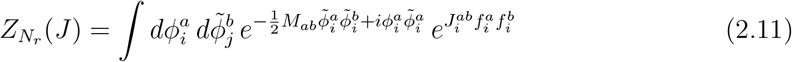

where we have defined

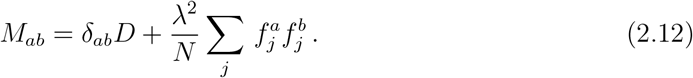

We see that the effect of the random variable *ξ*_*i*_ is to produce a quadratic term for 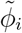 proportional to its variance *D*, while the Gaussian noise *W* introduces interactions with coupling *λ*^2^*/N*. When the activation function is well approximated by a linear regime, this gives a *ϕ*^4^-type interaction. In this work we will focus on more general nonlinear activation functions. This normalization with *N* is the natural one in order to produce finite correlation functions when *N* → ∞; see e.g. [21] for a review on the 1*/N* expansion. Next, the Gaussian 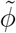 integral gives

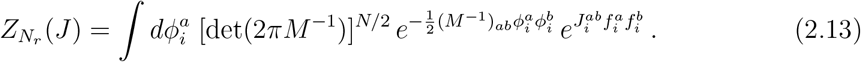

### 2.1 Collective fields

In order to perform a large *N* expansion we need to introduce collective fields so that the path integral becomes dominated by a “classical” saddle point. The model (2.13) is a particular case of a vector model, and the appropriate collective coordinate is [21]

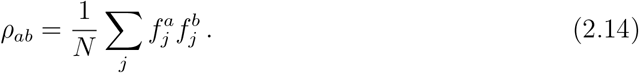

Note that this is a symmetric matrix in the replica indices. Introducing an auxiliary field *η*_*ab*_ that enforces this relation as a constraint^2^, we arrive at the following representation for the path integral:

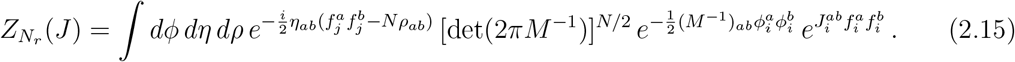

The key advantage of this representation is that now

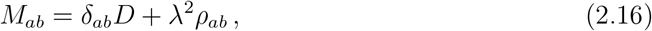

is independent of *ϕ*. The equivalence with (2.13) follows from the fact that the integral over *η* creates a delta function enforcing

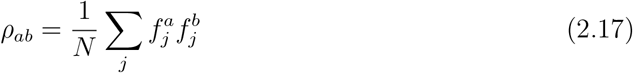

These fields will play the role of collective coordinates describing the correlations of the RNN.

The integrals appearing here cannot be obtained in general due to the nonlinear activation function. However, our main result will be that, when *N* ≫ 1, a semi-classical saddle point configuration controls the partition function. This will allow us to evaluate correlation functions analytically in a 1*/N* expansion, thus providing a new nonperturbative framework for understanding the dynamics of nonlinear RNNs. For related work on path integral representations with collective fields see also [23, 14].

## 3 Large *N* solution

The next step is to extract the quadratic dependence on *ϕ* from the activation function and then perform the *ϕ* integral. For this, we define a new function

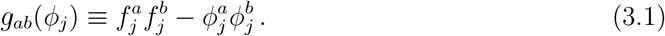

and so

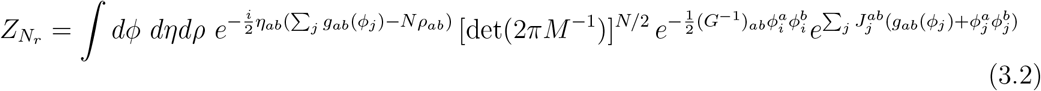

Where

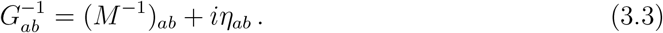

In the large *N* limit with the number of sources fixed, the contributions from the source terms are suppressed by 1*/N* compared to the other terms in (3.2). Therefore, in the limit of a large number of neurons, the highly complicated correlations of the neural system can be mapped into a dual classical configuration for the “master fields” *ρ* and *η*, which extremize the integrand of (3.2). The existence of such dual descriptions is one of the key advantages of large *N* systems, and has been very influential in the development of quantum field theory and gravity [24], and more recently also in deep learning [25]. Here we also find that it plays a central role in the understanding of nonlinear RNNs.

### 3.1 1*/N* and source expansion

At large *N*, the sources are then small perturbations of a network that is homogeneous, namely each neuron site *j* has the same dynamics. Let us denote a single neuron field *ϕ*^*a*^ by the variable *x*^*a*^, and introduce the normalized Gaussian average

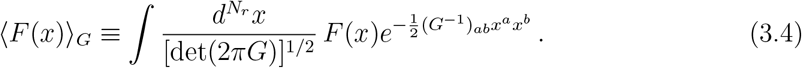

The partition function then becomes

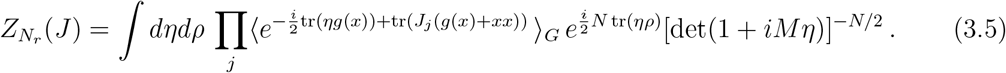

In order to shorten some of the following formulas, we will use a notation where *xx* is a matrix with elements (*xx*)^*ab*^ = *x*^*a*^*x*^*b*^, and 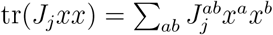.

We will consider an expansion for the exponential that resums the sources at each order.

For a single neuron index *j* we have

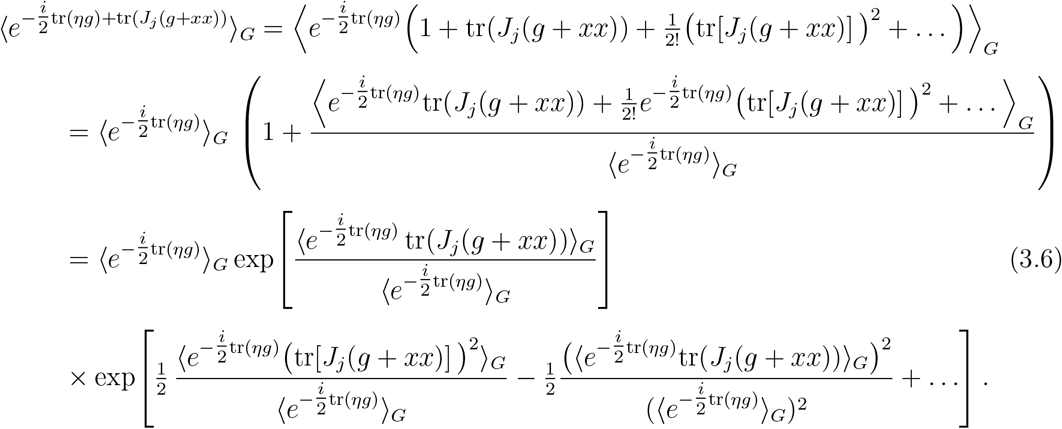

Similarly, the expectation value for the product over *j* indices becomes

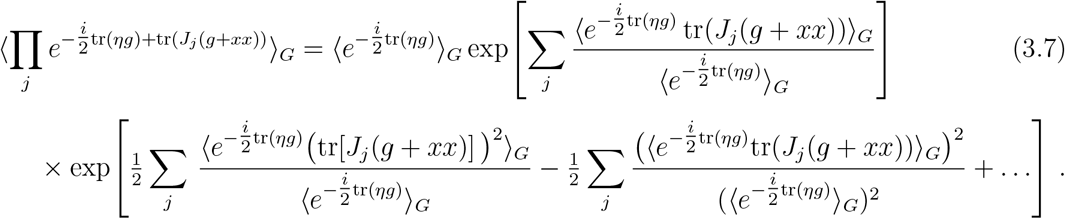

In the second line of this expression, we used the fact that contributions from different indices *j* factorize in the expectation value and hence cancel out between the two terms. By generalizing this procedure to higher orders we can compute the probability distribution for all neural correlators.

Here we will focus on calculating the partition function including second order contributions from the sources. We write Then

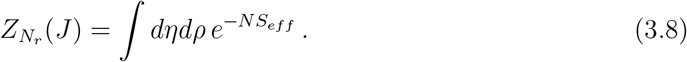

We will check self-consistently that the diagonal components of *ρ*_*ab*_ and *G*_*ab*_ are order *N* ^0^, while the off-diagonal components of *ρ*_*ab*_ and *G*_*ab*_, as well as all the variables *η*_*ab*_ are order 1*/N*. With this *N*-scaling, the leading terms at large *N* are

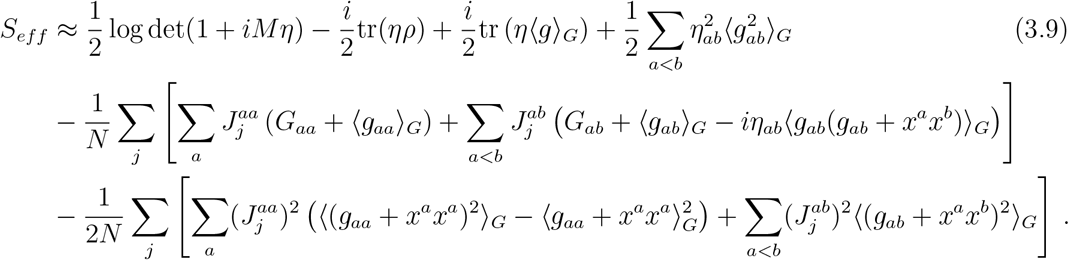

We have introduced the off-diagonal contribution 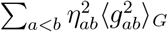 because it turns out to contribute at order 1*/N* ^2^; this will have the same *N*-scaling as *η*_*ab*_⟨*g*_*ab*_⟩_*G*_, which is the leading contribution for *a*≠ *b*. In contrast, the leading contribution from *η*_*ab*_⟨*g*_*ab*_⟩_*G*_ for *a* = *b*, occurs at order 1*/N*, while the corresponding quadratic term 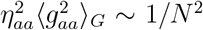 and hence can be neglected. The same *N*-scaling arguments have been used for the terms we kept in the other terms of (3.9).

### 3.2 Effects of nonlinearities

Let us now incorporate more explicitly the effects from the nonlinearity of the network, encoded in the insertions of *g*_*ab*_ in (3.9).

In order to evaluate the *G*-expectation values in the effective action, we will approximate *G*^−1^ in (3.4), taking into account that *G*_*aa*_ ∼ *N* ^0^ while *G*_*ab*_ ∼ *N*^−1^ if *b*≠ *a*. Then

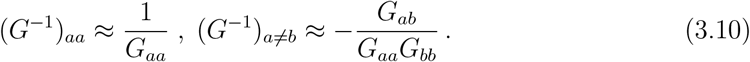

In these and the following expressions, we are not summing over repeated replica indices.

With this approximation,

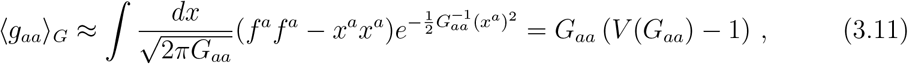

where we have defined

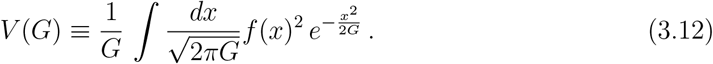

For the off-diagonal component, the leadning nonvanishing contribution comes from expanding the Gaussian exponent to first order in *G*_*a*≠*b*_,

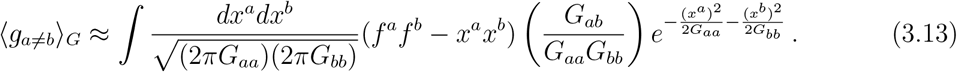

We introduce the function

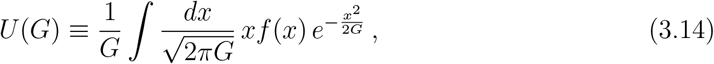

and then the off-diagonal nonlinearity contribution becomes

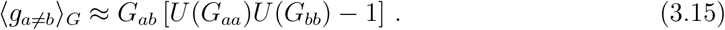

A similar calculation gives

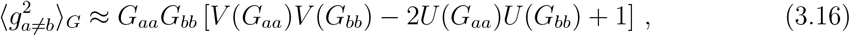

And

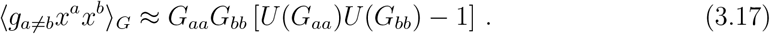

The last function we need to introduce comes from

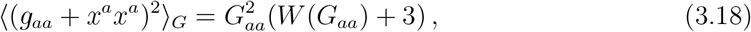

Namely

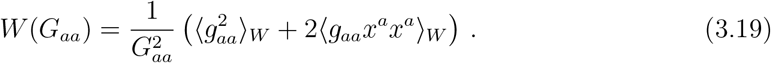

In this way, we arrive at our final form for the effective action,

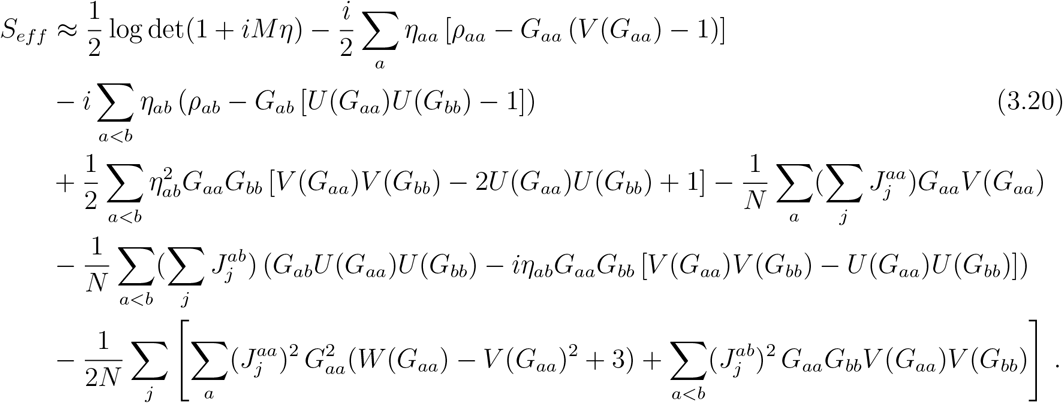

The functions *V, U* and *W* are Gaussian integrals containing the nonlinear activation function, while *M* and *G* are given in terms of *ρ* and *η* in Eqs. (2.16) and (3.3).

### 3.3 Saddle point description of neural correlation functions

At large *N*, (3.8) is dominated by the saddle point solution,

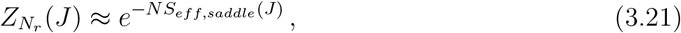

where the saddle point for the collective coordinates is determined by

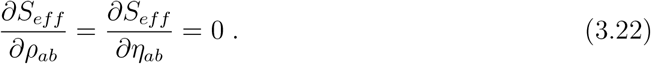

We will solve these equations in a 1*/N* expansion:

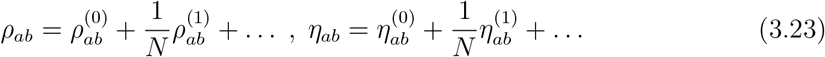

At order *N* ^0^, the result is

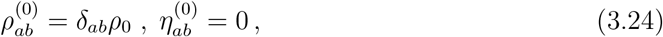

where *ρ*_0_ is the solution to

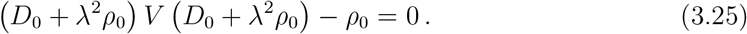

The fact that *η* = 0 at leading order provides a key simplification for the collective field description of nonlinear networks: it allows to perform a perturbative expansion of the interaction term ⟨ *e*^−(*i/*2)*ηg*^ ⟩_*G*_ in (3.5), as we did in (3.9).^3^ At this order, all the effects from the nonlinearities are then encoded in the self-consistent equation (3.25) for the collective field *ρ*. From (3.2) and (3.3), the two-point function for the input variables at order *N* ^0^ becomes

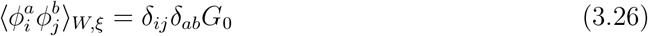

With

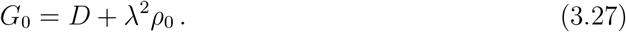

We will find it convenient to express our results in terms of this leading 2-point function *G*_0_. Replacing *ρ*_0_ in terms of *G*_0_, we find that the self-consistent equation (3.25) becomes

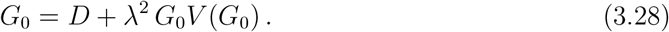

Recall that the function *V* (*x*) is determined in terms of the activation function via (3.12). We will give explicit examples below. In the cases of interest, this self-consistent equation shows that *G*_0_(*λ, D*) is a monotonic function of the coupling *λ*. It is then simpler to just use (3.28) to write *λ*^2^ as a function of *G*_0_, and this is what we use for the analytic results in the next section.

It is also useful to rewrite this self-consistent equation in terms of

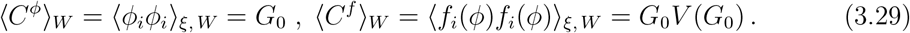

Then (3.28) becomes

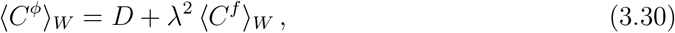

which we recognize as a mean field equation. Such expressions have appeared before, e.g. in [27, 23]. It agrees with the self-consistent equation (2.2) found very recently in [18].

For the linear network, *f* (*x*) = *x, V* (*G*) = 1, and (3.28) can be solved to give

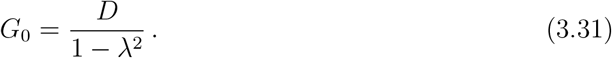

Therefore, the 2-point function diverges as the coupling *λ* → 1. This limit corresponds to the instability of the linear network, where the linear restoring term in (2.1) is overcome by the effects from fluctuations of th connectivity matrix. Understanding the network dynamics beyond this point requires including effects beyond the linear approximation. In our present formalism, we are including nonlinear effects exactly in the 1*/N* approximation; this divergence is resolved for activation functions that are bounded by *f* (*x*) *< x*^*p*^ for *p <* 1 at large *x*. To see this, Eq. (3.28) can be solved at large *G*_0_ approximating the integral in *V* (*G*_0_) by the regime where *f* (*x*) ∼ *x*^*p*^; the self-consistent solution then shows that the growth of *G*_0_ is bounded by *G*_0_ ≲ (const) *λ*^2/(1−*p*)^. See Sec 4.1 for more details.

At the next order in the 1*/N* expansion, the collective fields 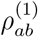 and 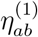 are proportional to linear and quadratic combinations of the sources. In other words, at order 1*/N* the collective fields encode the backreaction of the insertions of sources and the interplay with the network nonlinearity. Let us exhibit these effects explicitly at linear order in the sources:

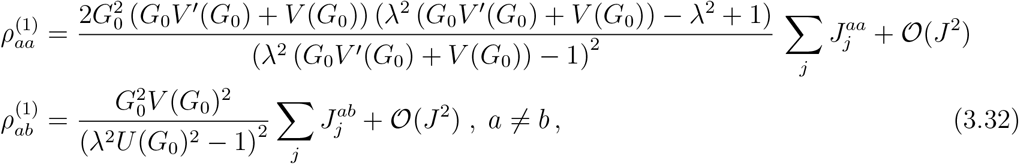

And

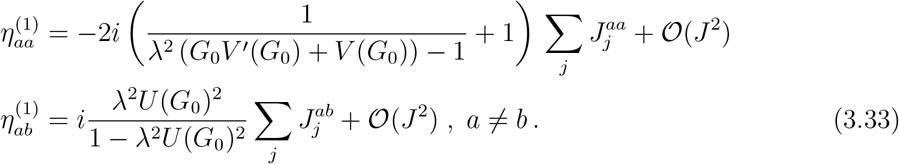

The quadratic terms are straightforward to compute, we won’t use their explicit expression. Finally, computing the 1*/N* saddle point values 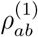 and 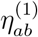 replacing into the effective action, we obtain the generating function of connected correlation functions,

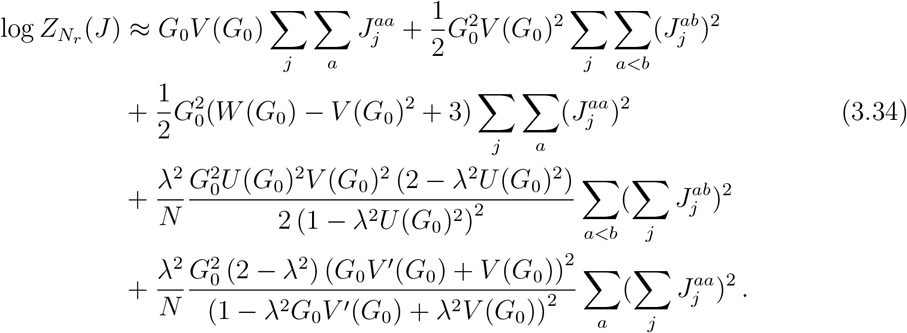

The first two lines in this expression have been simplified by using the self-consistent equation (3.28). The third and fourth lines can also be written in different ways by applying the self-consistent equation. For our results below we will use the form where *λ*^2^ is solved for in terms of *G*_0_ using (3.28). In particular, this will be useful for showing that 1 − *λ*^2^*U* (*G*_0_)^2^ is strictly positive, and can only vanish for a linear activation function.

Let us present these results in terms of neural correlation functions. We define the covariance matrix

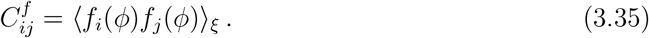

Then the output 2-point function at large *N* becomes

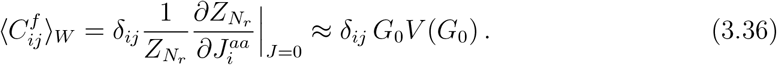

From (3.12), this is the Gaussian expectation value of *f* (*x*)^2^, as expected. Let us now consider the 2-point functions of the covariance matrix *C*^*f*^. The first case is

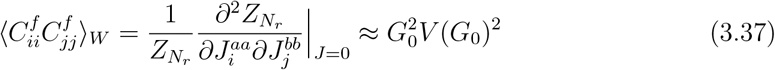

where *i* ≠ *j* are some neuron indices, and *a* ≠ *b* are some replica indices. Therefore, these fluctuations approximately factorize at large *N*,

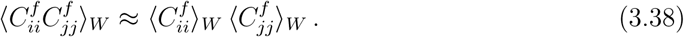

We also find factorization if *i* = *j*:

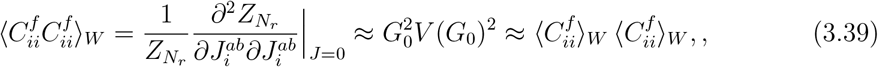

where again *a* ≠ *b* are two different replica indices. The same result is obtained by differentiating with respect to 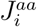 and 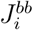.

On the other hand, the fluctuations of off-diagonal elements of the covariance matrix are

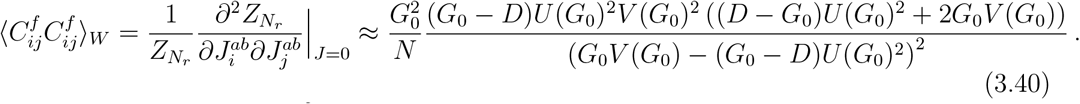

Here we have eliminated *λ*^2^ in terms of the self-consistent equation. Let us use this form to prove that 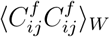 cannot diverge as long as *f* (*x*) is bounded by *x*^*p*^, *p <* 1, at large *x*. We already saw that *G*_0_ is finite in such a nonlinear network, so this correlation function can diverge only if the denominator vanishes. The terms inside the square in the denominator are

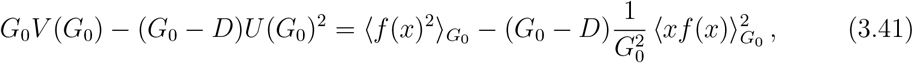

where 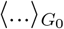 denotes the normalized Gaussian average (3.4) but now with respect to a single variable *x* with variance *G*_0_. By the Cauchy-Schwarz inequality,

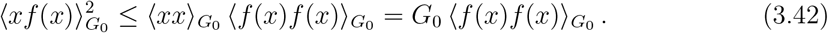

Therefore,

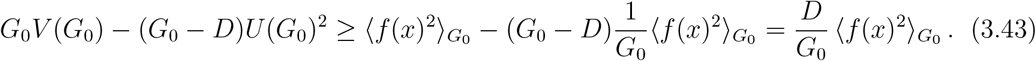

The right hand side is strictly positive, and we conclude that in the presence of interactions *f* (*x*) bounded by *x*^*p*^ with *p <* 1 at large *x*, the correlation of off-diagonal components of the covariance matrix is finite at all coupling.

This analysis also yields the statistics of higher correlation functions. For instance, at this order of the large *N* expansion

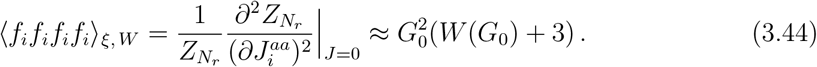

A general property of the large *N* expansion is that, once a leading nonzero correlator is identified, higher point correlation functions factorize. Two examples are shown in (3.38) and (3.39). Similarly, having identified the leading off-diagonal contributions (3.40), higher-point correlators including it will factorize. For instance,

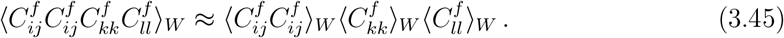

An important consequence of this property is that we now have a complete characterization of the probability distribution *P* (*C*^*f*^) of the covariance matrix 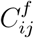: its generating functional is given by a subset of the terms in (3.34),

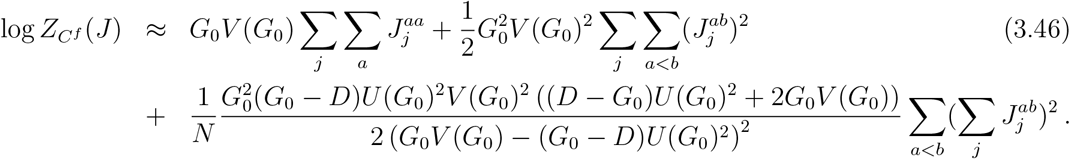

In other words, the statistics of *C*^*f*^ is determined by the leading nonzero correlators (3.36), (3.40), plus large *N* factorization.

### 3.4 Dimension of participation

Computations in neural systems usually depend on structured neural activity whose dimensionality is lower than the one of the state space [12, 28], that would be *N* for our system. One of the most usual ways to quantify dimensionality is the *participation dimension*. It is derived from the eigenspectrum of the neuronal covariance matrix. This matrix underlies Principal Component Analysis [29], and indicates how pairs of neurons covary across time and task parameters [12]. It is defined by

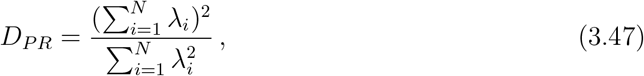

where *λ*_*i*_ are the eigenvalues of the covariance matrix. The motivation for this definition is that if the activity has predominant variability along a set of *D* directions, then there will be *D* eigenvalues that are similar between them but much larger than the rest and we will have *D*_*PR*_ ≈ D.

Rewriting this expression in terms of the trace of the covariance matrix and its square we obtain that

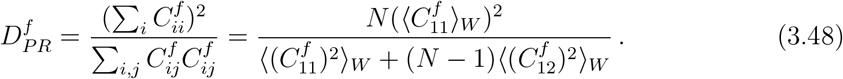

We used the self-averaging property to relate the sum over neurons to the *W*-average, and the homogeneity of the network to fix the sites at 1 and 2. At large *N*, and recalling that 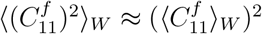, this becomes

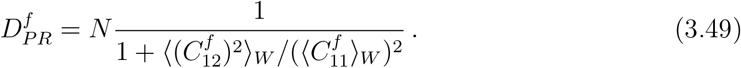

Using the results of the previous subsection, we obtain

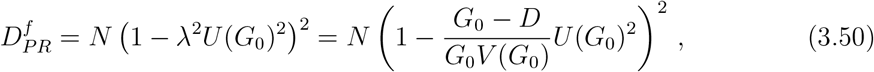

where in the second equality we have eliminated the coupling using the self-consistent equation. By the same arguments as before in terms of the Cauchy-Schwarz inequality, the participation dimension is strictly positive for activation functions bounded by *x*^*p*^ with *p <* 1.

All these steps can be redone with minor modifications in order to compute the statistics of the correlation functions of inputs *ϕ*_*i*_; see the Appendix A for more details. In particular, the participation dimension for inputs becomes, from (A.10),

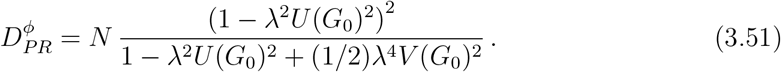

Therefore, except in the linear network, the effective dimension for outputs and inputs is different. We will evaluate explicitly these correlation functions for different networks in the next section.

### 3.5 Comparison to other works

Closely related references to our work include [23, 14], which also used a path integral representation and the large *N* limit. Ref. [23] considered the nonlinear case and focused on the auto-covariance and time-dependent effects, but did not include 1*/N* effects. They obtained a self-consistent equation which plays the role of our (3.28). Similar self-consistent equations also appear in other references such as [27, 16].

On the other hand, Ref. [14] focused on the linear network. In this work a path-integral formulation is also used. In contrast to our work there are no replicas and a linear source term is used. Correlations up to order four are evaluated taking succesive derivatives with respect to the source and the mean values and variances of the elements of the covariance matrix are determined using a Wick decomposition of the temporal averages. This is possible in the linear case because the statistics for fixed couplings is Gaussian. This approach is not possible in the non-linear case, requiring the use of replicas.

The recent preprint [18] proposed an ansatz to solve the nonlinear network in terms of an effective coupling. This is motivated by mean field theory considerations and is verified numerically and using random matrix theory. This ansatz can be derived explicitly from our large *N* treatment. From (3.50), the role of the effective coupling is played by

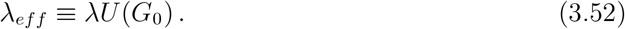

To verify that this gives the proposal of [18], we observe that integrating by parts in Eq. (3.14), we obtain

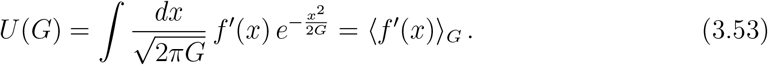

The dimension of participation (3.50) becomes

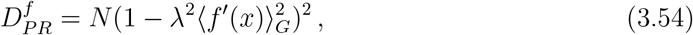

which coincides with Eq. (3.1) of [18] at vanishing frequency. Our result for ⟨ *C*_*ij*_*C*_*ij*_ ⟩_*W*_ is also consistent with theirs.

## 4 Applications

In this section we will consider applications of our general results above. We will first consider a class of power-law activation functions for which we obtain explicit analytic expressions for the correlation functions. They are characterized by power-law dependence on the coupling *λ*, and are useful for approximating more realistic networks in different regimes, such as near saturation, weak coupling, etc. We next introduce a class of “Padé activation functions” that are very useful for simultaneously capturing the small *ϕ* and large *ϕ* regimes of the activation functions. With this choice we will also be able to give explicit results for the correlation functions (albeit in terms of special functions). We will then perform extensive comparisons with numerical results for two activation functions of interest, and we will assess the behavior of 1*/N* corrections and the approach to equilibrium.

### 4.1 Power-law activations

As our first application, we will obtain analytic results for activation functions of the form

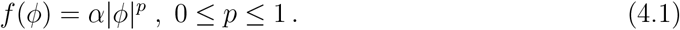

Such power-law activation functions appear in different contexts. The exponent *p* = 1 corresponds to the linear activation function; this is also a good approximation for odd activation functions and sufficiently small *ϕ*. The case *p* → 0 applies to a regime near saturation, such as large *ϕ* for the th(*x*) activation. Other powers, such as *p* = 1/2 can be relevant over broad intermediate regimes for *ϕ*. In fact *p* = 1/2 is obtained for neural dynamics that undergo a type I bifurcation that leads to spike generation [30, 31, 32]. For our present analysis, we will assume that the neural activation function admits a range of parameters where it can be well approximated by (4.1).

The functions *V* (*G*) and *U* (*G*) introduced in (3.12) and (3.14), respectively, evaluate to

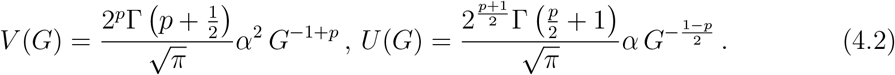

The self-consistent equation for the 2-point function ⟨*ϕ*_*i*_*ϕ*_*j*_⟩_*ξ,W*_ = *G*_0_ becomes, from (3.28),

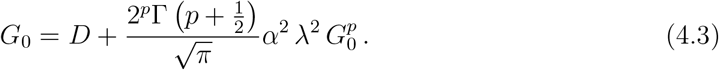

For *p* = 1, this reproduces the standard 2-point function for linear networks,

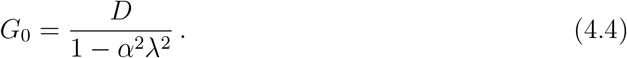

*α*^2^*λ*^2^ → 1 corresponds to the instability of the linear network. This instability is avoided by the interactions created by the nonlinear activation function.

On the other hand, taking *p* → 0 we see that the 2-point function for networks in the saturation regime is

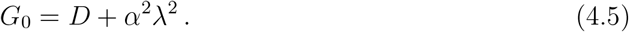

For more general *p*, (4.3) can be solved in a power series of *λ*^2^ for small *λ*. This is the weakly coupled limit of the interacting network. The self-consistent equation also allows us to obtain the 2-point function in the limit of strong coupling (large *λ* or small *D*). The result is

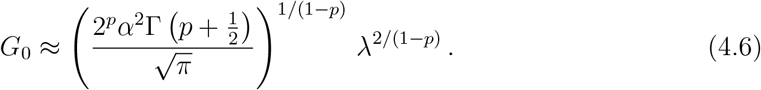

The 2-point correlation of the outputs then scales as

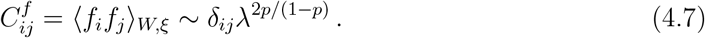

This shows concretely how any sub-linear transfer function is enough to suppress the divergence that appears in linear systems.This is compatible with the result one would obtain for the analysis of a rate model in the mean field limit [33].

Plugging the explicit expressions for *V* and *U* in (3.40), we obtain the variance of the off-diagonal elements of the covariance matrix:

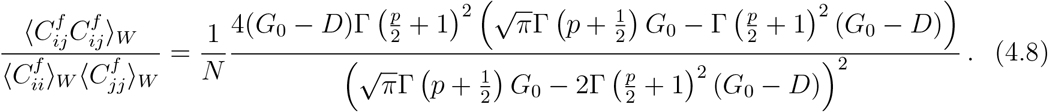

In the three regimes that we discussed above, this expression simplifies to

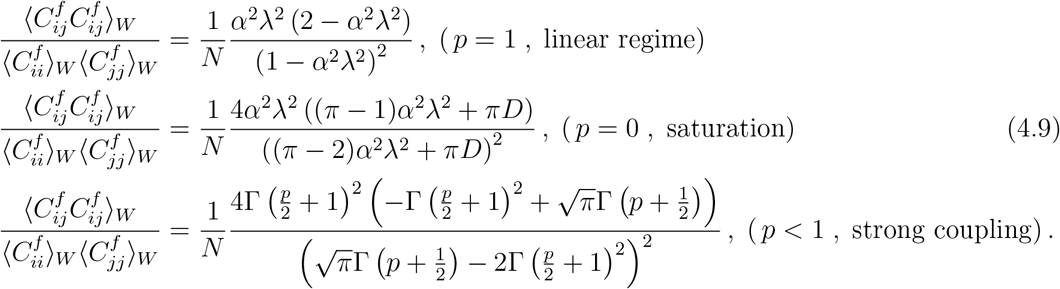

We can also evaluate explicitly the participation dimension. Using (4.8), we obtain

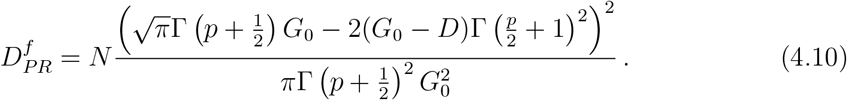

In the three regimes we are discussing, the effective participation dimension evaluates to

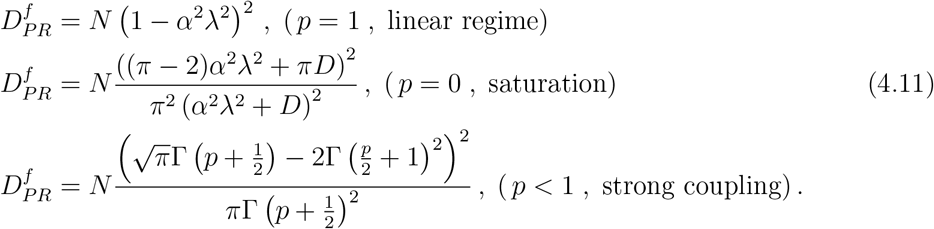

The result for the participation dimension of the linear network has been obtained before in [19, 15]; the effective dimension vanishes as *α*^2^*λ*^2^ → 1, corresponding to the onset of the instability of the linear network. On the other hand, for nonlinear networks, with *p <* 1, this function does not vanish. The effective dimension is monotonically decreasing with *p*, attaining its maximum at *p* = 0 (saturation regime), and vanishing for *p* = 1 at the instability threshold.

### 4.2 Padé activations and numerics

In this section we will perform detailed comparisons between our large *N* formulas and numerical results. The details of the numerical methods are presented in Appendix B.

We introduce a class of nonlinear transfer functions that we call “Padé activation functions”:

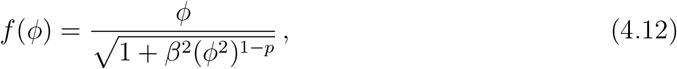

where the parameter *β* controls the strength of the nonlinear behavior. One motivation for this is that many neural systems can be modeled by an activation function that is linear at small input *ϕ*, but exhibits saturation at large input, or a power-law behavior over an intermediate but large regime. The Taylor expansion approximation can in general capture either the limit of small or large *ϕ*, but not both. The idea of Padé approximants is that they can capture both regimes simultaneously by performing an approximation in terms of ratios of polynomials [34]. This type of activation function has been previously analyzed in the context of convolutional neural networks [35] and deep learning [36] but not in RNNs, as far as we know. This expansion allows to perform analytically the Gaussian integrals such as Eqs. (3.12) and (3.14). This motivated our choice of (4.12): its square is a ratio of polynomials with the desired small and large *ϕ* behavior. In particular, for *p* = 0 the square of (4.12) is the Padé approximant of (th(*βϕ*)*/β*)^2^. In what follows we will focus on two cases: *p* = 0 and *p* = 1/2.

#### 4.2.1 Activation function with p = 0

Let us first evaluate our results and compare with numerics for an activation function

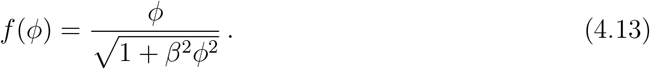

This form, unlike the th(*x*) function, allows us to evaluate explicitly the functions *V* (*G*) and *U* (*G*) that enter the neural correlators. We find

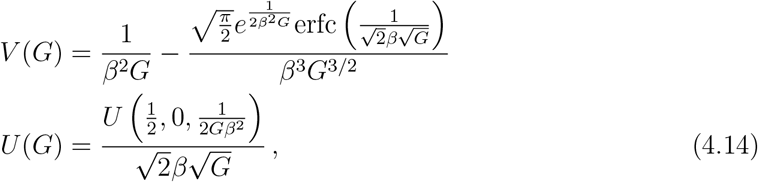

where erfc(*x*) is the complementary error function, and *U* (*a, b, z*) is the confluent hypergeometric function.

We now compare these results with the ones obtained using network simulations (see Appendix B for details). In Fig. 1 we show the average values of the diagonal and nondiagonal terms of the covariance matrix and the participation dimension of the outputs.

**Figure 1.**
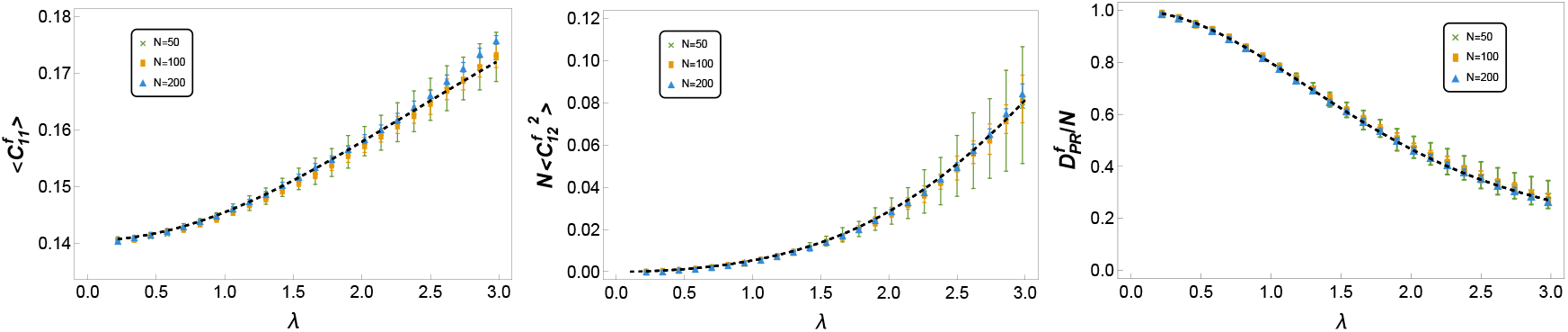
Comparison of analytical (dashed line) and numerical results (dots) for the output correlations. Left panel: average value of the diagonal correlation. Central panel: average of the square of the off-diagonal correlators. Right panel: participation ratio (participation dimension divided by network size). We use the transfer function of Eq.(4.13) with *β* = 2 and *D* = 1; the network sizes are *N* = 50, 100, 200. The error bars represent the standard deviation over 5 simulations with different realizations of the coupling matrix *W* and noise *ξ*.

We find an excellent agreement between theory and simulation even for sizes of about a few hundred neurons. In Fig. 1 it is possible to see that the standard deviation of the different quantities decrease with network size. This is compatible with the 1*/N* expansion presented in Section 3.1. The fluctuations are controlled by higher order moments that fall with *N* with 1*/N* ^2^ or faster. In order to check this we performed simulations with sizes up to *N* = 800 and evaluated the rate of decrease of the standard deviation of the average diagonal and off-diagonal terms of the output correlations. The results are shown on Fig. 2.

**Figure 2.**
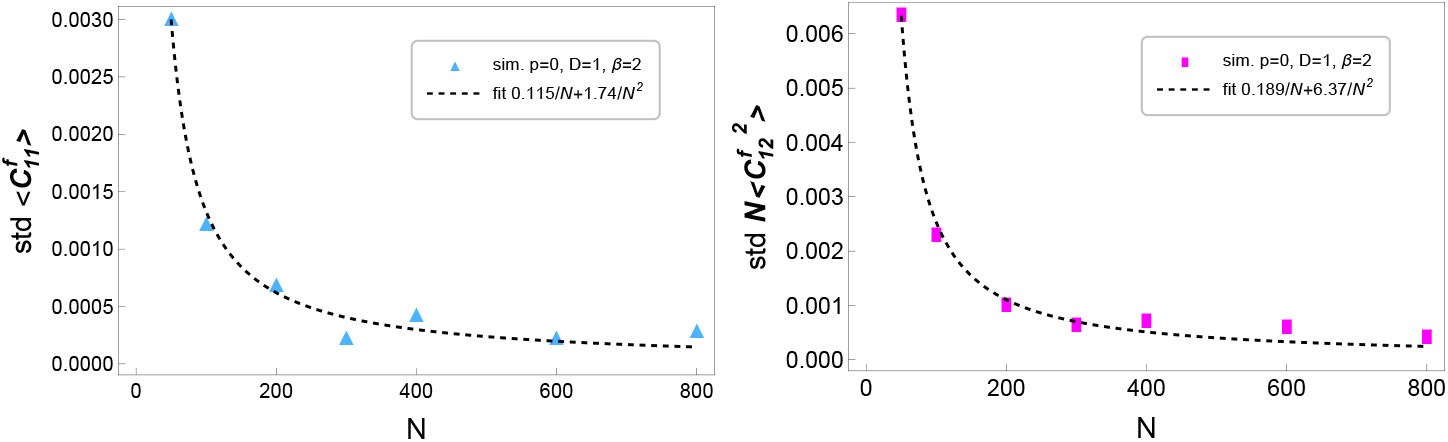
Scaling with *N* of the standard deviations of 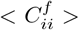 (left panel) and 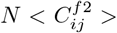 (right panel). Dashed line shows a fit in powers of 1*/N* : *a/N* + *b/N* ^2^.

In order to test the robustness of the results with respect to the simulation time we also performed simulations with a fixed number of steps *n*_*t*_ (that corresponds to a simulation time *t*_*sim*_ = *n*_*t*_δ) without enforcing the condition *d <* 10^−4^. These results are shown in Fig. 3. We find thte results converge between *n*_*t*_ = 100 and 200 iterations. This represents a simulation time between 10 and 20 times the characteristic time constant *τ*.

**Figure 3.**
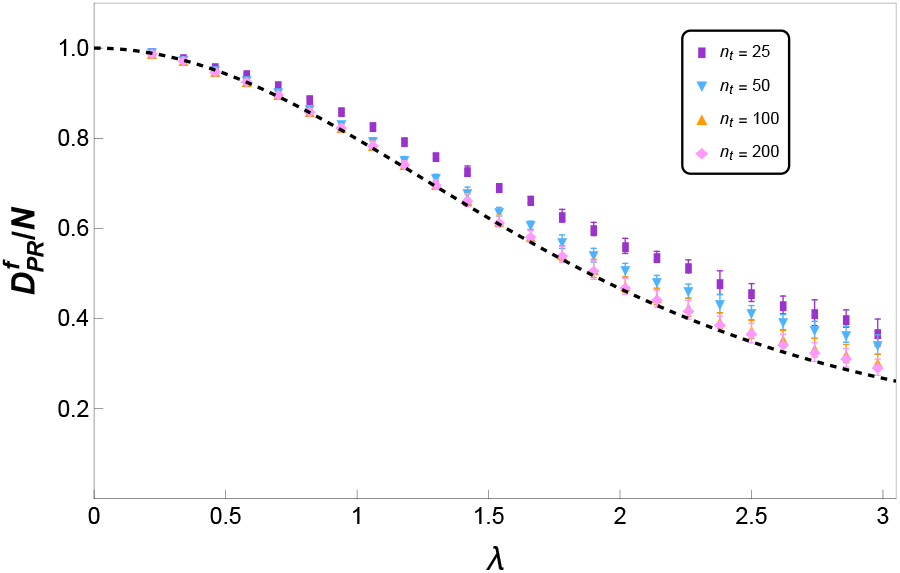
Participation ratio for different simulation times. Dashed line: analytical result. Saturating transfer function (*p* = 0), *D* = 1 and *β* = 2.

#### 4.2.2 Activation function with p = 1/2

Next we consider,

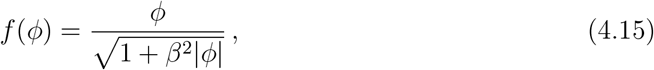

| which at large *ϕ* gives a power-law behavior *f* ∼ | *ϕ* | ^1/2^.

The functions *V* and *U* are slightly more involved but can still be computed explicitly in terms of special functions,

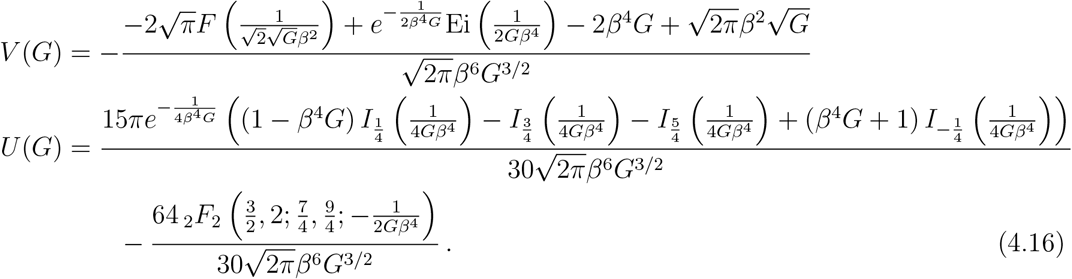

Here *F* (*z*) is the Dawson integral, Ei(*z*) is the exponential integral function, *I*_ν_(*z*) is the modified Bessel function of the first kind, and _*p*_*F*_*q*_(*a*; *b*; *z*) is the generalized hypergeometric function.

In Fig. 4 we show the results for the non-saturating Pade function of Eq. (4.15). We again obtain an excellent agreement between the analytic formulas and the numerical results.

**Figure 4.**
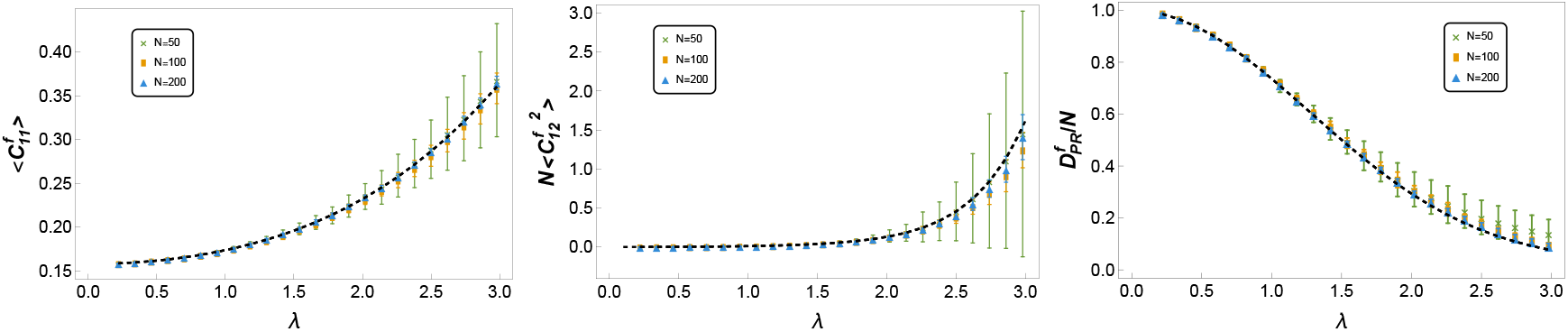
Comparison of analytical (dashed line) and numerical results (dots) for the output correlation, for the Padé activation function with *p* = 1/2. Left panel: average value of the diagonal correlation. Central panel: average off-diagonal correlators. Right panel: participation ratio (participation dimension divided by network size). We use the transfer function of Eq. 4.15 with *β* = 2 and *D* = 1. The network sizes are *N* = 50, 100, 200. The error bars represent the standard deviation over 5 simulations with different realizations of the coupling matrix *W* and noise *ξ*.

## 5 Conclusions

In this work we have developed a path-integral framework to compute the statistics of correlations in nonlinear recurrent networks, including 1*/N* effects that are required to evaluate cross-correlations of the covariance matrix and the participaton dimension. Our approach generalizes previous treatments of linear networks by incorporating arbitrary nonlinear activation functions as interaction terms in the effective action. This resolves the instability of the linear theory, yielding strictly positive participation dimensions and a rich set of correlation structures.

A central outcome of our analysis is the emergence of a small set of collective variables that capture the full statistics of correlations. This representation makes explicit the way in which global network dynamics constrain local pairwise correlations. It also highlights the universality of some of the results: for instance, the 1*/N* suppression of cross-correlations persists in the nonlinear regime, but their relative fluctuations become essential in controlling the participation dimension. This is consistent with previous findings in the linear case [14, 15], while substantially extending their range of applicability. We also obtained the generating function of connected neural correlators, focusing for the most part on the covariance matrix; see Sec. 3.3. This is another key result of our work, and it is based on large *N* factorization.

The explicit results obtained for power-law and the new class of Padé activation functions illustrate two complementary aspects of the theory. Power-law activations exhibit scaling behavior controlled by the strength of the recurrent coupling, revealing how changes in network connectivity can reshape correlation statistics. Padé activations, on the other hand, provide a flexible family of transfer functions that remain tractable analytically while capturing important features of realistic nonlinearities. The agreement between our analytic predictions and numerical simulations across these cases gives strong support to the validity of the approach. Numerical simulations indicate that the results of the large *N* limit are relevant even for sizes of a few hundred neurons, while the time required to perform the averages is not required to be larger than about 20 synaptic time constants.

Our results also allow us to relate the path-integral method to alternative theoretical frameworks. Dynamical mean-field theory and random matrix methods, which have been successfully applied to related questions [16, 17, 18], have provided valuable insights into the statistics of covariance matrices. The path-integral formulation complements these approaches by emphasizing the role of collective variables, the effects of external sources, and the description in terms of the generating function, which naturally encode higher-order correlators and enable systematic expansion in 1*/N*. This complementary perspective could prove useful in situations where non-Gaussian features of the activity, or finite-size effects, play an essential role.

More broadly, the ability to derive analytic predictions for correlation statistics in nonlinear recurrent networks opens the door to several applications. In neuroscience, it provides a theoretical tool to interpret experimental measurements of correlations, dimensionality, and variability in cortical recordings. In machine learning, the connection between correlation structure and participation dimension may shed light on representational capacity in large recurrent architectures. Finally, from a methodological standpoint, the path-integral formalism can be adapted to other complex systems where nonlinear interactions and collective dynamics shape correlation statistics.

In future work, it will be important to extend our approach to characterize out-of-equilibrium correlation functions, where temporal structure and nonstationary dynamics are expected to play a crucial role. Furthermore, the path-integral formulation provides an efficient framework to evaluate subleading 1*/N* effects using standard diagrammatic methods. For the recurrent networks studied here, such corrections originate from Gaussian fluctuations around the nontrivial saddle point as well as from interaction vertices induced by nonlinear activation functions. The path-integral framework is also promising for incorporating additional structures, such as plasticity. Exploring these contributions will deepen the connection between neural dynamics and field-theoretic techniques, and we expect to report on these developments in future work.

## Acknowledgments

GM is supported by CONICET (PIP grant 112202101 00114CO), CNEA and Instituto Balseiro, Universidad Nacional de Cuyo, FR is supported by a CONICET PhD fellowship and by Instituto Balseiro, Universidad Nacional de Cuyo. GT is supported by CONICET (PIP grant 11220200101008CO), CNEA, Instituto Balseiro, Universidad Nacional de Cuyo, and a Simons Foundation targeted grant to institutions.

### A Correlations of neural inputs

In this Appendix we discuss briefly how to obtain correlations of neural inputs,

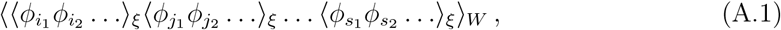

This requires modifying (2.6) so that the source couples to *ϕ*_*i*_*ϕ*_*j*_ instead of *f*_*i*_*f*_*j*_. The steps are the same as in Secs. 2 and 3, with the replacement 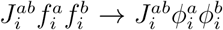. The net effect is that (3.2) is modified to

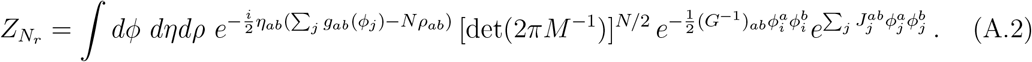

The large *N* effective action (3.9) is then replaced by

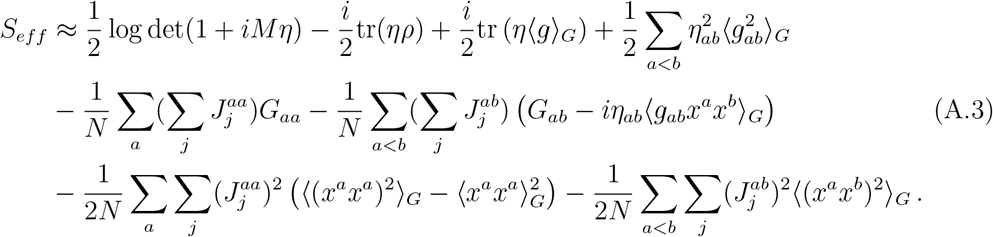

Recalling the parametrization of Sec. 3.2, this becomes

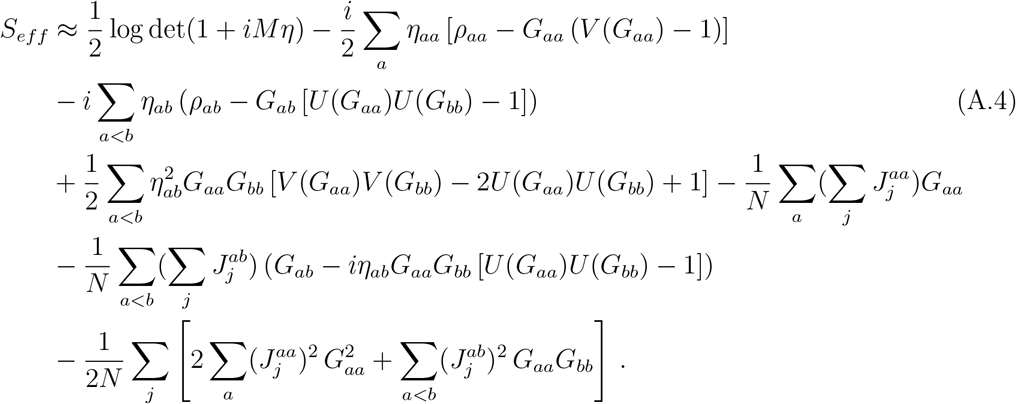

Computing the saddle point for *ρ* and *η* in a 1*/N* expansion and plugging back into the effective action, we arrive at the generating function for connected correlators,

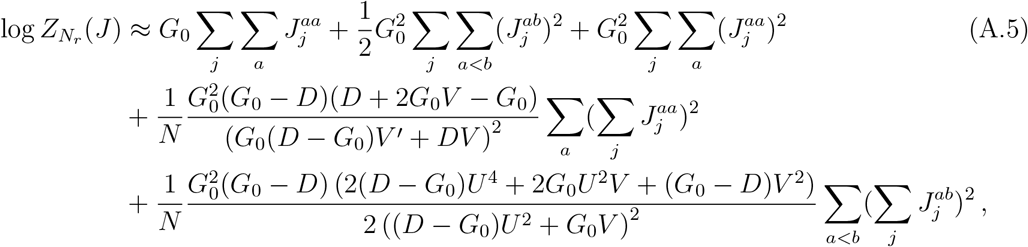

which should be compared with the analog expression (3.34) for the outputs. Here *U* = *U* (*G*_0_) and *V* = *V* (*G*_0_) are the functions defined in the main text, and *V* ^*′*^ = *V* ^*′*^(*G*_0_).

At leading order in the large *N* expansion, the two-point function is

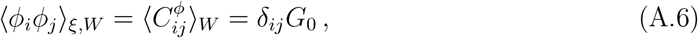

and *G*_0_ satisfies the self-consistent equation (3.28),

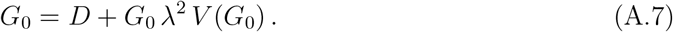

Diagonal correlators of the *ϕ* covariance matrix factorize,

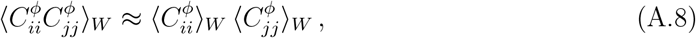

while the off-diagonal correlators are given by

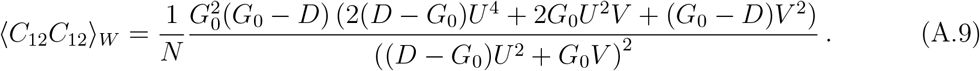

This predicts a participation dimension for inputs

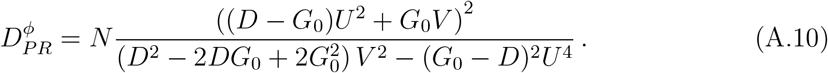

We have written all these expressions by eliminating *λ*^2^ in terms of the self-consistent equation. In the main text, we give an equivalent expression for the participation dimension in terms of *λ*^2^, see (3.51).

In Fig. 5 we show the comparison between input and output participation ratios. We see two different behaviors: for the high noise case *D >* 1, we find that *D*_*PR*_ decreases with *λ* and the gap between the input and output dimensions is very small. In contrast, in the low case of low noise we find an increasing *D*_*PR*_ and a significant dimension gap, in agreement with [16].

**Figure 5.**
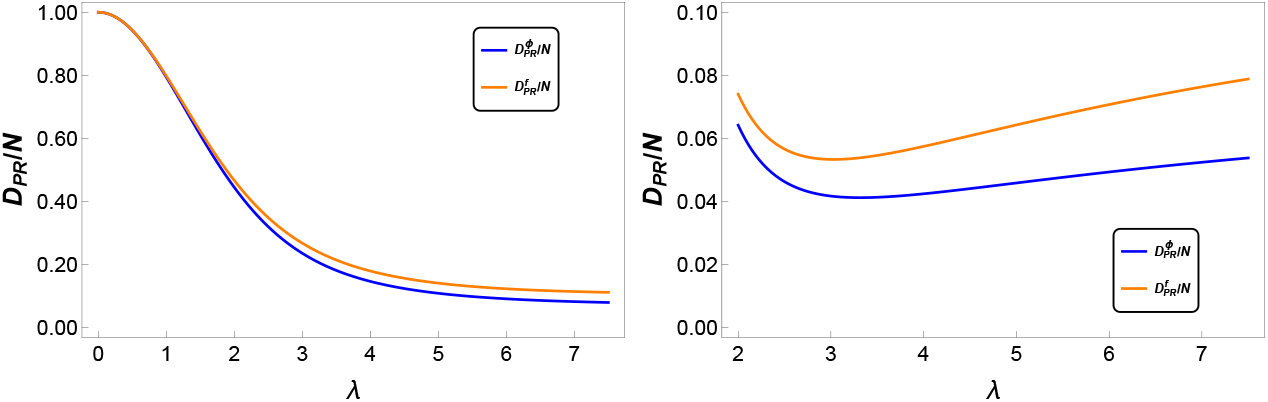
Comparison of participation ratios of inputs and outputs, for *β* = 2, *D* = 1 (left) and *β* = 2, *D* = 0.1, right. Note the difference in scales in the y-axis between the plots.

The comparison between the analytical results and numerical simulations for the input correlations are shown in Figs. 6 and 7. As in the case of output correlations we fond an excellent agreement even for not very large network sizes.

**Figure 6.**
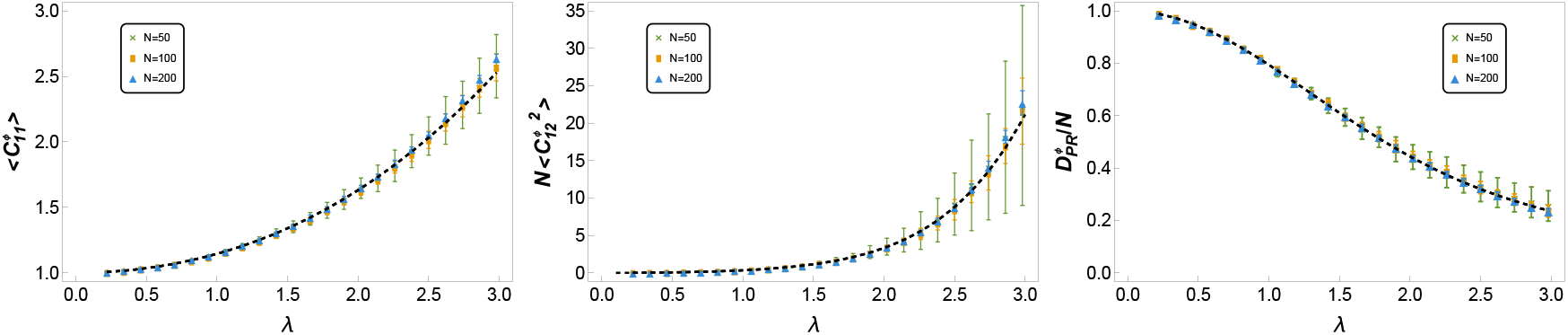
Comparison of analytical and numerical results for the input correlation. Left panel: average value of the diagonal correlation. Central panel: average off-diagonal correlators. Right panel: participation ratio (participation dimension divided by network size). We use the transfer function of Eq.(4.13) with *β* = 2 and *D* = 1. Network sizes *N* = 50, 100, 200. The error bars represent the standard deviation over 5 simulations with different realizations of the coupling matrix *W* and noise *ξ*.

**Figure 7.**
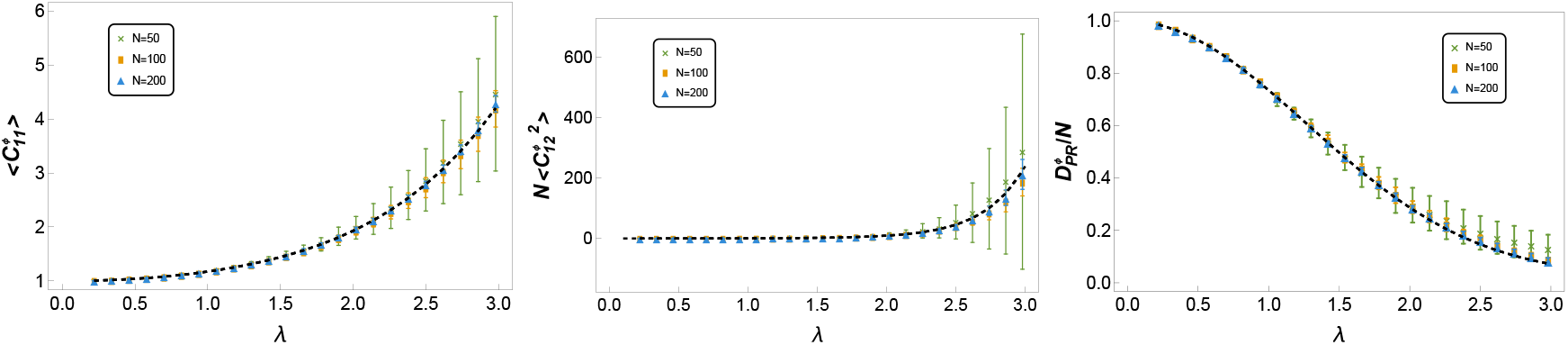
Same as in the the previous figure but with the non saturating activation function of Eq. (4.15).

### B Network simulations

The differential equations that control the network dynamics, Eq.(2.1), are simulated using a first order Euler method. As we intend to be in the limit *τ* → 0 at each time step we choose a vector of random Gaussian variables *ξ*_*i*_ (1 = 1, …*N*) with zero mean and variance *D* and iterate

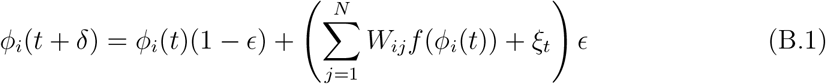

until the mean quadratic difference 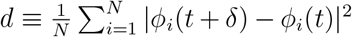 is smaller than 10^−4^. We use δ = 0.01, *ϵ* = 0.1. This procedure is repeated *n*_*l*_ = 1000 times to obtain the covariance matrices:

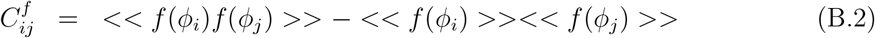

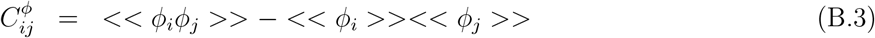

where the brackets denote the temporal average: 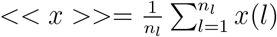 generates bias in the statistics, specifically in the second order terms we implemented the corrections proposed in [19] (Supplementary Material).

## Footnotes

1 We could also choose it to be symmetric in the indices (*a, b*), at the cost of introducing factors of 2 between the source derivatives of *Z*(*J*) and the correlation functions.

2 *η*_*ab*_ is also symmetric in (*a, b*).

3 A similar simplification was observed for linear networks in [14].

